# Cell type resolved co-expression networks of core clock genes in brain development

**DOI:** 10.1101/2020.12.30.424790

**Authors:** Surbhi Sharma, Asgar Hussain Ansari, Soundhar Ramasamy

## Abstract

The circadian clock regulates vital cellular processes by adjusting the physiology of the organism to daily changes in the environment. Rhythmic transcription of core Clock Genes (CGs) and their targets regulate these processes at the cellular level. Circadian clock disruption has been observed in people with neurodegenerative disorders like Alzheimer’s and Parkinson’s. Also, ablation of CGs during development has been shown to affect neurogenesis in both in vivo and in vitro models. Previous studies on the function of CGs in the brain have used knock-out models of a few CGs. However, a complete catalog of CGs in different cell types of the developing brain is not available and it is also tedious to obtain. Recent advancements in single-cell RNA sequencing (scRNA-seq) has revealed novel cell types and elusive dynamic cell states of the developing brain. In this study by using publicly available single-cell transcriptome datasets we systematically explored CGs-coexpressing networks (CGs-CNs) during embryonic and adult neurogenesis. Our meta-analysis reveals CGs-CNs in human embryonic radial glia, neurons and also in lesser studied non-neuronal cell types of the developing brain.

## Introduction

Self-sustained oscillatory expression of a core set of genes is known to drive circadian rhythms in organisms ranging from fungi and archaebacteria to humans. At the molecular level, these rhythms arise from the waxing and waning of transcription factors which gives rise to rhythmicity in gene expression of their targets. In recent years, several groups have shown the association of key components of the molecular clock with neuronal function and diseases. Specifically, CLOCK protein, a part of the positive arm of the transcription-translation feedback loop has been shown to affect migration of neuronal cells (Fontenot et al. 2017). Mutations in CGs have been associated with various disorders; single nucleotide changes in NR1D1, PER1 and NPAS2 have been shown to be linked with the etiology of autism spectrum disorders (Goto et al. 2017; Nicholas et al. 2007). The reduced expression of CLOCK has been observed in epileptogenic tissues in humans (P. Li et al. 2017). Also, knock-out of BMAL1 the binding partner of CLOCK, in mice has been shown to lower the seizure threshold (Gerstner et al. 2014). Circadian disruption has also been observed in major depressive disorder (MDD), as the rhythmicity of CGs like BMAL1, PER1-2-3, NR1D1, BHLHE40-41 and DBP was disrupted in the postmortem brains derived from MDD patients (J. Z. Li et al. 2013).

The brain development and function is maintained in large part by embryonic and adult neurogenesis. During cortical development, heterogeneous populations of neural stems cells are present in the subventricular zone (SVZ) and ventricular zone (VZ), while differentiating neurons form cortical plate (CP). Previous studies have shown the involvement of CGs in neurogenesis; BMAL1 and PER2 are involved in cell cycle regulation in adult neurogenesis (Bouchard-Cannon et al. 2013). Also, BMAL1 knockout mice exhibit symptoms of ageing and perturbed ocular parameters (Yang et al. 2016). Similarly, conditional mutant mice of Clock or Bmal1 show delayed maturation of inhibitory parvalbumin neurons in the visual cortex (Kobayashi, Ye, and Hensch 2015).

Above studies clearly establish the role of CGs in the brain development, through genetic manipulation of selected CGs. Systematically extending the above strategies to all CGs is both laborious and time consuming. In case of the developing human brain it is further complicated due to ethical reasons. In this study we analyzed scRNA-seq datasets of embryonic and adult neurogenic niches to comprehensively explore the cell type identity and expression dynamics of CGs in neurogenesis using pseudotime analysis. Since most of the CGs function as transcription factors (TFs) with a wide range of genomic targets (Koike et al. 2012), we also identified their co-expressing networks (CNs). For pseudotime and co-expression analysis we used TSCAN and SCENIC respectively (Figure1 & refer methods). Our analysis reveals CGs-CNs enrichment in distinct cell types of the neurogenic niches thereby providing framework for future studies on their role in neurodevelopmental disorders. In addition to CGs-CNs we also provide CNs of all transcription factors as a resource for future investigations.

**Figure 1:**
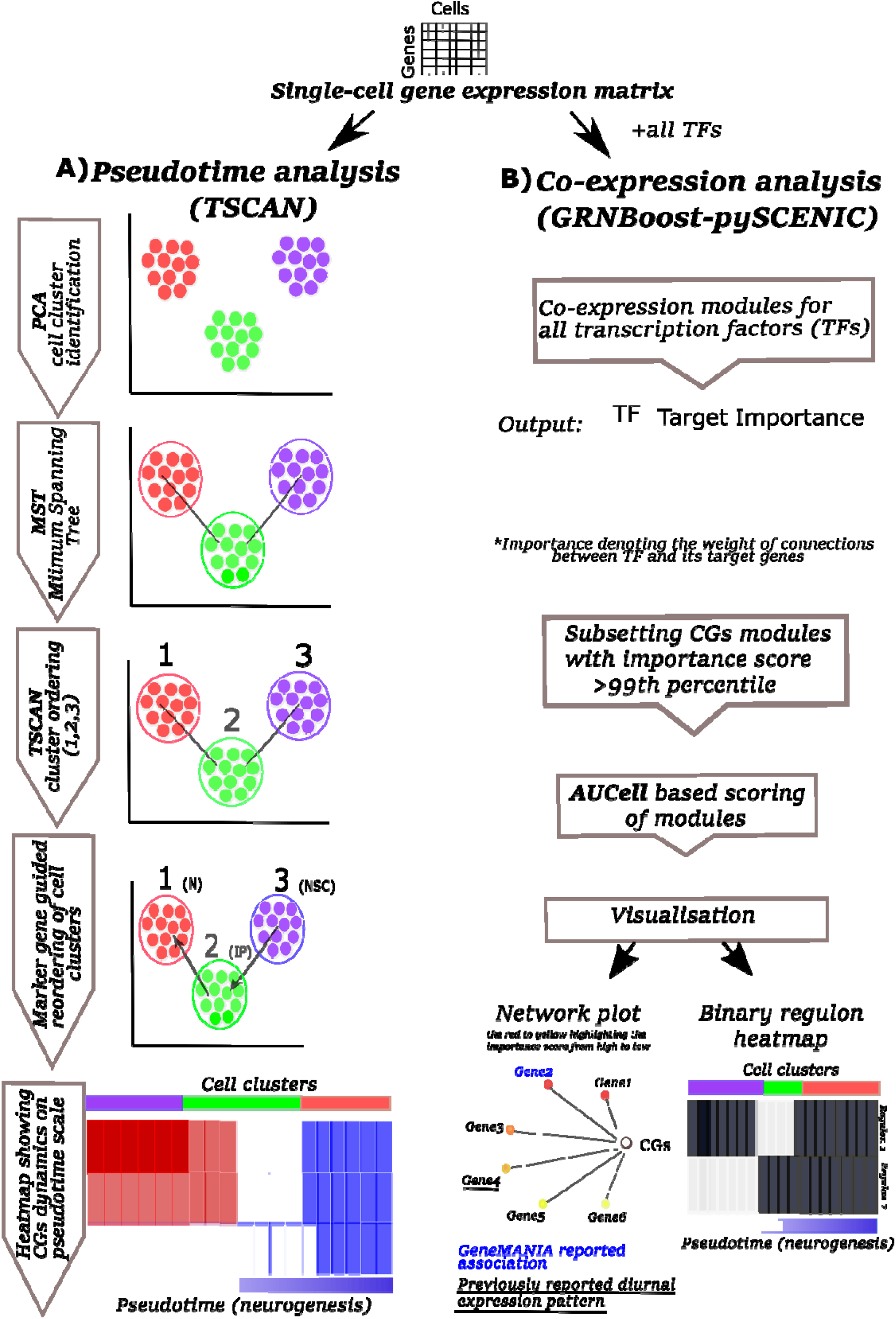
Schematic depicting workflow of A) pseudotime & B) co-expression analysis using TSCAN and GRNBoost respectively.

## Results

CGs that from the focus of this study are listed in supplementary Table S1.

### Expression dynamics of CGs-CNs in mouse embryonic cortex E15.5

Embryonic neurogenesis is characterized by the rapid proliferation of neural stem cells. These cells divide symmetrically and expand the pool of neurogenic stem cells during the initial stages of neurogenesis. Later stages of embryonic neurogenesis are characterized by asymmetric divisions to give rise to neurons via intermediate progenitor cells (Götz and Huttner 2005). In contrast to embryonic neurogenesis, adult neurogenesis is restricted to specialized regions like SVZ and SGZ (Petrik and Encinas 2019). Apart from giving rise to neurons and glia, embryonic neural stem cells also serve as the source of adult neural stem cells. We intended to analyze the expression of CGs with cell type identity and expression dynamics in mouse embryonic and adult neurogenesis.

To study the CGs expression dynamics during embryonic neurogenesis we have used a singlecell transcriptome dataset of mice generated by Yuzwa et. al., which encompass embryonic timepoints: E11.5, E13.5, E15.5 and E17.5 days (Yuzwa et al. 2017). Using TSCAN (Ji and Ji 2016), neural progenitors and differentiating neuron clusters were identified from the above datasets. Pax6 and Tubb3 marker genes were used to identify neural progenitors and differentiating neurons respectively. Further these cells were pseudotime aligned with neural progenitors at the start and differentiating neurons at the end. Upon this neurogenesis trajectory aligned by pseudotime, statistical significance of expression dynamics of CGs was calculated. Out of all the developmental time points studied, only E15.5 showed statistically significant expression dynamics of three CGs – Rorb, Nfil3 and Csnk1e (Figure 2A). Except for Csnk1e, the two CGs expressions were high along the pseudotime associated with differentiated neurons.

**Figure 2:**
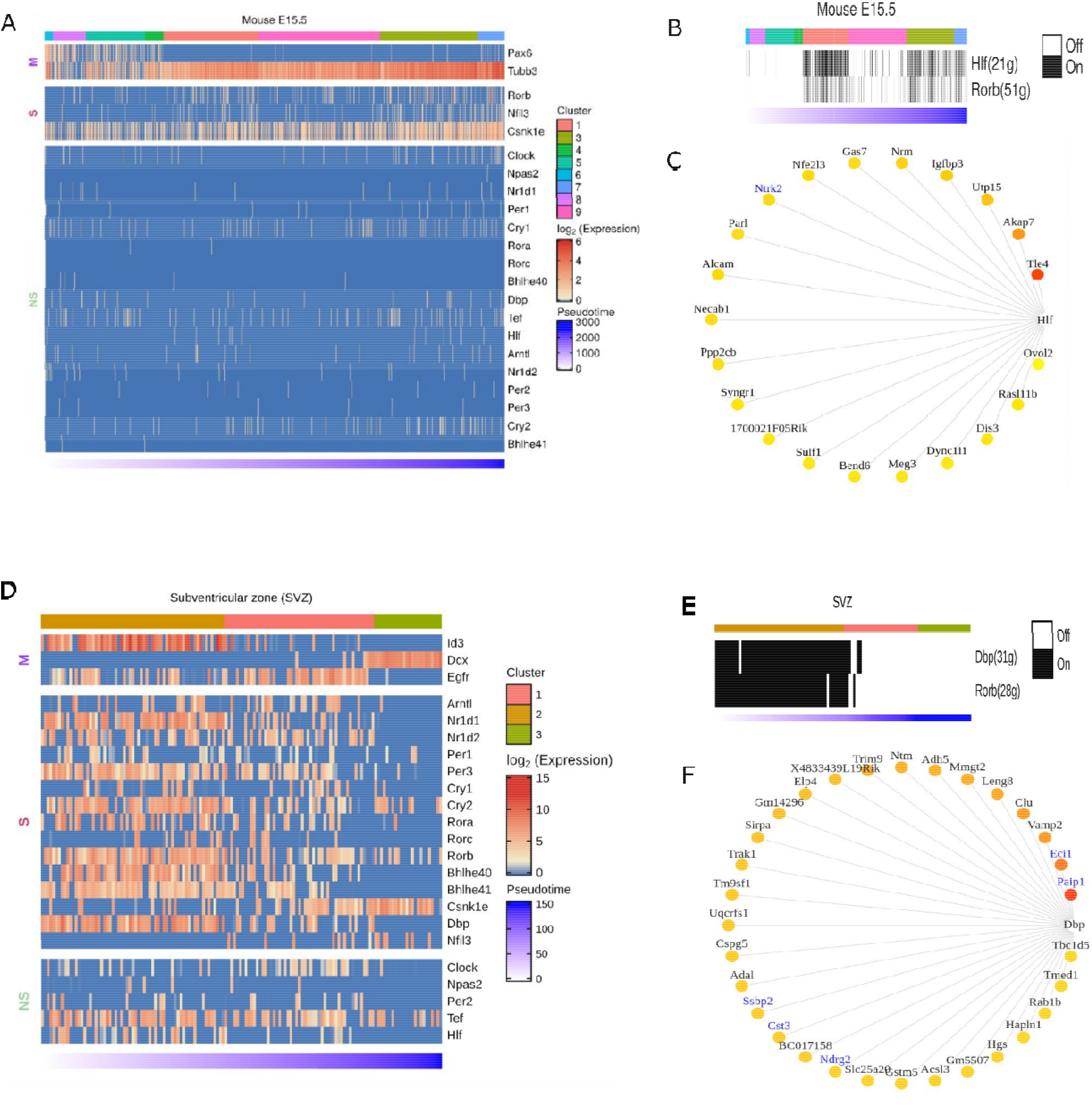
Expression dynamics of CGs-CNs in mouse embryonic and adult neurogenesis. A&D) Heatmap showing CGs (row) expression over pseudotime ordered cells (column) in neurogenesis trajectory of A) mouse E15.5 cortex, D) mouse sub ventricular zone (SVZ), top annotation denotes cell clusters assigned by TSCAN, bottom annotation denotes TSCAN calculated pseudotime in ascending order. In heatmaps, M denoting marker genes, S and NS denoting statistically significant and non-significant pseudotime expression dynamics respectively. B&E) Binary activity heatmap of CG modules over pseudotime aligned neurogenesis trajectory of B) mouse E15.5 cortex E) mouse SVZ. Top annotation denotes cell clusters assigned by TSCAN, bottom annotation denotes TSCAN calculated pseudotime in ascending order. C&F) Network plot showing CGs and its target genes. The edges of the target genes with highest importance score (red) to lowest (yellow). GeneMANIA reported co-expression gene pairs are indicated in blue color.

Applying the co-expression analysis workflow on E15.5 dataset revealed two gene modules: Hlf(21g) and Rorb(51g) (Figure 2B). Hitzler JK et al earlier hypothesized the role of Hlf in neuronal differentiation based on its expression pattern in diverse regions of developing mouse brain (Hitzler et al. 1999), also experimental models of epilepsy show downregulation of Hlf (Rambousek et al. 2020). Rorb, shows expression in the L4 region of the mouse and human brain (Zeng et al. 2012). It also plays a vital role in the development and organization of the whisker barrel in rodent brain (Clark et al. 2020). These studies provide hints for involvement of Hlf and Rorb in the brain development, additionally our analysis discloses their underlying gene regulatory network (Figure 2C & S1). Out of 21 genes in the Hlf module (Figure 2C), Ntrk2 was previously reported in the GeneMANIA co-expression database.

### Expression dynamics of CGs-CNs in the neurogenic niches of adult mouse brain

In the adult brain, neurogenesis occurs in restricted niches: sub granular zone (SGZ) of the hippocampus and sub ventricular zone (SVZ) near the lateral ventricles (Doetsch, García-Verdugo, and Alvarez-Buylla 1999; Kriegstein and Alvarez-Buylla 2009). The periodic activation of qNSCs in SGZ/SVZ produces mature neurons migrating to deep layers of dentate gyrus (DG) and olfactory bulb respectively, whose homeostasis plays a vital role in learning, memory formation and emotion regulation (van Praag, Kempermann, and Gage 1999; Toda et al. 2019). SGZ and SVZ regions are shown to have expressions of CGs (Bouchard-Cannon et al. 2013). Using publicly available scRNA-seq data and pseudotime analysis, we studied the CGs expression dynamics during adult neurogenesis.

For pseudotime ordering of SVZ (Llorens-Bobadilla et al. 2015) we used known marker genes like Id3 and Egfr for quiescent and activated NSCs respectively and Dcx for differentiated neurons (Figure 2D). Most of the CGs analyzed showed significant enrichment in initial stages of pseudotime which corresponds to the transition from quiescent to activated neural stem cells. Though most of the CGs analyzed showed enrichment during early stages of adult neurogenesis, Csnk1e showed reverse trend of enrichment, with higher expression in the later stages of pseudotime corresponding to the birth of differentiated neurons.

Besides SVZ, we also analyzed SGZ, the other prominent adult neurogenic niche using scRNA-seq data (Shin et al. 2015). It also showed enrichment of CGs expression during early stages of neurogenesis (Figure S2), though the expression dynamics was not as evident as in SVZ. Both SVZ and SGZ showed enrichment of Csnk1e during later stages of adult neurogenesis.

Our co-expression analysis on SVZ and SGZ, revealed CGs-CNs only in SVZ datasets as none of the predicted CNs of SGZ datasets passed the AUCell scoring threshold. In SVZ, the coexpression analysis revealed two distinct CGs-CNs: Dbp(31g) and Rorb(28g) active during early stages of adult neurogenesis (Figure 2E). The detailed analysis of the Dbp module (Figure 2F) revealed 5 of its target genes (Paip1, Eci1, Ssbp2, Cst3 and Ndrg2) to be documented in the GeneMANIA. Among the above genes, Ndrg2 expression was previously reported in SVZ (L. Liu et al. 2012).

### Expression dynamics of CGs-CNs in developing human brain

The development of human brain and maturation lasts upto to 20 years (Peña-Melian 2000), while embryonic and fetal developmental changes range from 0-8 and 8-24 post-conception week (pcw) respectively (Silbereis et al. 2016). To identify CGs-CNs in early human brain development, we analyzed the scRNA-seq dataset generated by Nowakowski et al. (Nowakowski et al. 2017). The dataset includes prefrontal cortex (PFC) and primary visual cortex (V1), with ages ranging from 5 to 37pcw. In total, from 48 samples they broadly identified 11 major cell types. Their dataset is also available as an interactive web browser (https://cells.ucsc.edu). We applied co-expression analysis workflow on their dataset and used the cell cluster information provided by authors.

Our co-expression analysis revealed four CGs-CNs: BHLHE41(193g), BHLHE40(57g). RORB(12g) and NR1D2(14g) (Figure 3A).

**Figure3:**
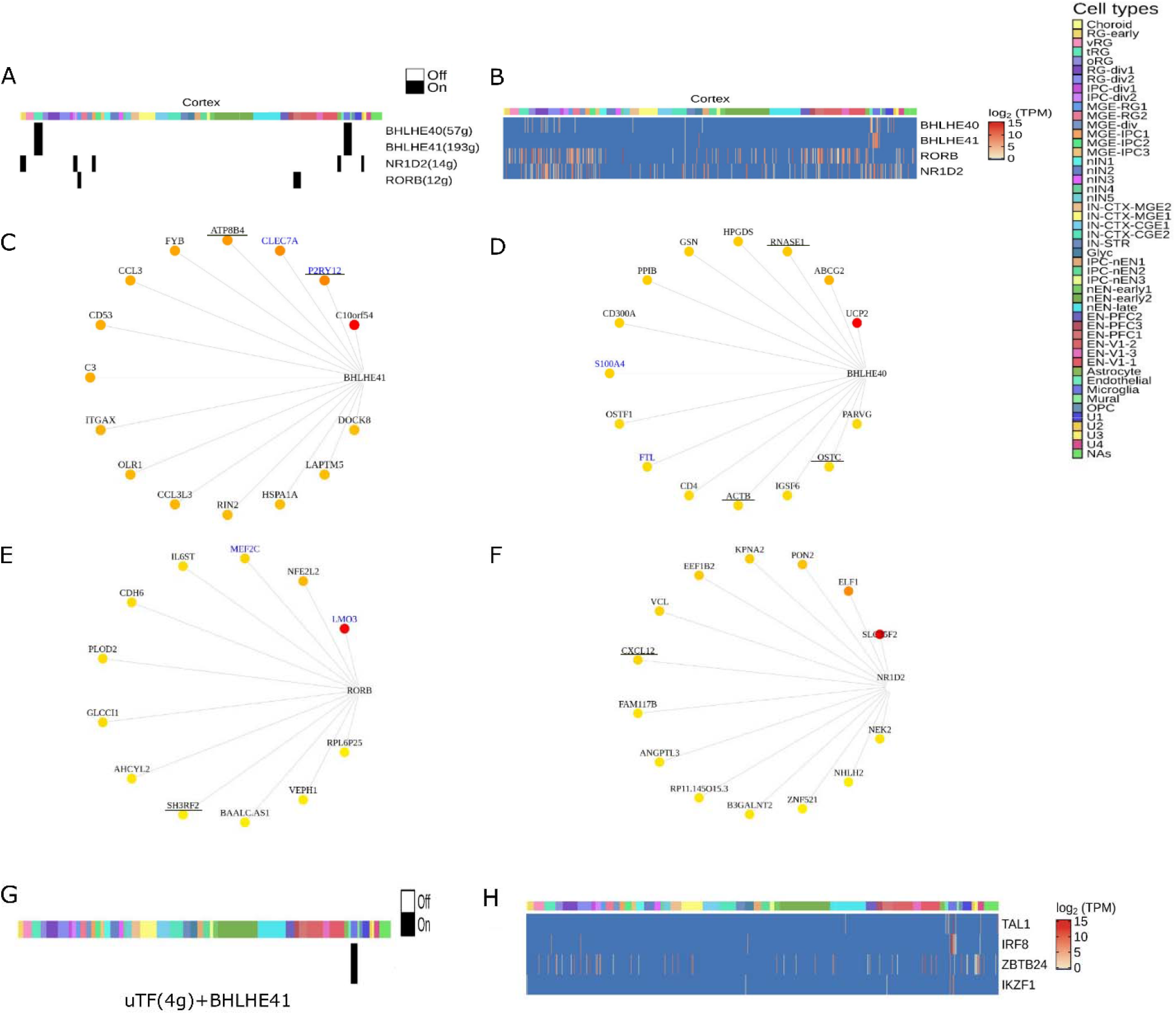
Expression dynamics of CGs-CNs in developing human brain. A) Binary activity heatmap of CGs modules over cell clusters of the developing human brain, cell clusters as annotated by Nowakowski et al. for cluster description refer table S4. B) Heatmap showing gene expression of CGs over cell clusters, which showed significant enrichment score in the co-expression analysis. Cell clusters as annotated by Nowakowski et al. for cluster description refer table S4. C, D, E & F) Network plots showing CGs and its targets in the developing human brain, the edges are placed from highest (red) to lowest (yellow) importance score of target genes. GeneMANIA reported co-expression gene pairs are highlighted in blue color, only top15 target genes in the BHLHE40/41 module are shown, target genes with known diurnal expression are underlined. G) Binary activity heatmap of upstream TFs co-expressing with BHLHE41 over cell clusters of the developing human brain, cell clusters as annotated by Nowakowski et al. for cluster description refer table S4. H) Heatmap showing gene expression of upstream TFs co-expressing with BHLHE41, which showed significant enrichment score in the co-expression analysis. Cell clusters as annotated by Nowakowski et al. for cluster description refer table S4.

Interestingly, the BHLHE41(193g)/BHLHE40(57g) module enriched predominantly in nonneuronal cell types like mural, microglial and truncated radial glia (tRG) cluster (Figure 3A). Few topmost genes co-expressing with BHLHE41 (Figure 3C) included P2RY12, CLEC7A, ATP8B4 and FYB. P2RY12 functions as a receptor for nucleotides which are released in response to CNS (Haynes et al. 2006) and blood brain barrier injury (Lou et al. 2016) to assist chemotaxis of microglia to the site of injury. CLEC7A has been shown to be linked with beta amyloid disease progression (Wirz et al. 2013). Genes including CLEC7A, P2RY12 and FYB serve as marker of microglia and we reveal their co-expression with BHLHE41 in our analysis. In BHLHE40(57g) module (Figure 3D), S100A4 was one of the co-expressing target genes, with reported co-expresssion in GeneMANIA. It also acts as a biomarker of glioblastoma (Chow et al. 2017). Among the top connected genes in the same module we observed HPGDS, which is known to localize to microglia and upregulated in Alzheimer’s (Mohri et al. 2007).

The RORB(12g) module (Figure 3E) showed enrichment in medial ganglionic eminence-radial glia cluster 1 (MGE-RG1) and excitatory neurons-PFC 1 (EN-PFC1) cluster, the RORB module was also captured in the developing cortex and SVZ dataset of mouse. Comparison of the target genes of the RORB module in mouse and humans revealed MEF2C-RORB gene pair coexpression in mouse and human embryonic cortical neurogenesis. Both genes are known to play a vital role in corticogenesis (H. Li et al. 2008), with mutations in MEF2C causing mental retardation disorder (Zweier et al. 2010).

NR1D2(14g) expression was enriched in radial glia early (RG-early), intermediate progenitor cells (IPC-div2), medial ganglionic eminence-intermediate progenitor cells (MGE-IPC1) and astrocytes (Figure 3A). Knockdown of NR1D2 in adult murine neural stem cells has been shown to affect neural differentiation (Shimozaki 2018). The topmost target genes of the NR1D2 module included SLC35F2, ELF1, PON2, KPNA2 and EEF1B2 (Figure 3F). These genes are involved in vital neuronal functions like SLC35F2 in transport across blood brain barrier (Mochizuki et al. 2020), ELF1 in neurite growth (Gao et al. 1996) and PON2 imparts neuroprotection by scavenging superoxides (Costa et al. 2014).

The enrichment of four CGs-CNs in diverse cell types of the developing human brain prompted us to look for upstream regulators of CGs. Our search for upstream TFs co-expressing with CGs revealed only one significant upstream module of BHLHE41 [uTF(4g)+BHLHE41] (Figure 3G). We found four TFs namely TAL1, IRF8, IKZF1 and ZBTB24 which are documented to have regulatory role in microglia activation in response to injury (Masuda et al. 2012), aging (Wehrspaun, Haerty, and Ponting 2015) with TAL1 acting as a specific marker of human microglia (Galatro et al. 2017).

### Expression dynamics of CGs-CNs during astrocyte genesis

Previously, considered to be the support system of neurons in the brain, glial cells are being recognized as vital players in brain function. By promoting neuronal survival and modulating synapse function these cells actively participate in neuronal signal transduction. In particular, participation of astrocytes in circadian regulation has been recognized recently (Tso et al. 2017). The various functions of astrocytes like modulation of intracellular calcium levels, neurotransmitter release occur rhythmically. A study by (Lananna et al. 2018) has shown that astrocyte specific genetic deletion of Bmal1 and behavioral disruption of rhythms in mice induces astrocyte activation. This effect is augmented when Bmal1 is knocked out in both neurons and astrocytes. These studies point to an important role for astrocytes in maintaining circadian function in the brain.

We analyzed the total transcriptome of human fetal and adult brain derived astrocytes (Ye Zhang et al. 2016). In contrast to neurons, astrocytes showed an upregulation of CGs namely CLOCK, CRY2, PER3, RORA, RORB, HLF, NR1D1, BHLHE40 and BHLHE41 upon maturation (Figure 4A).

**Figure 4:**
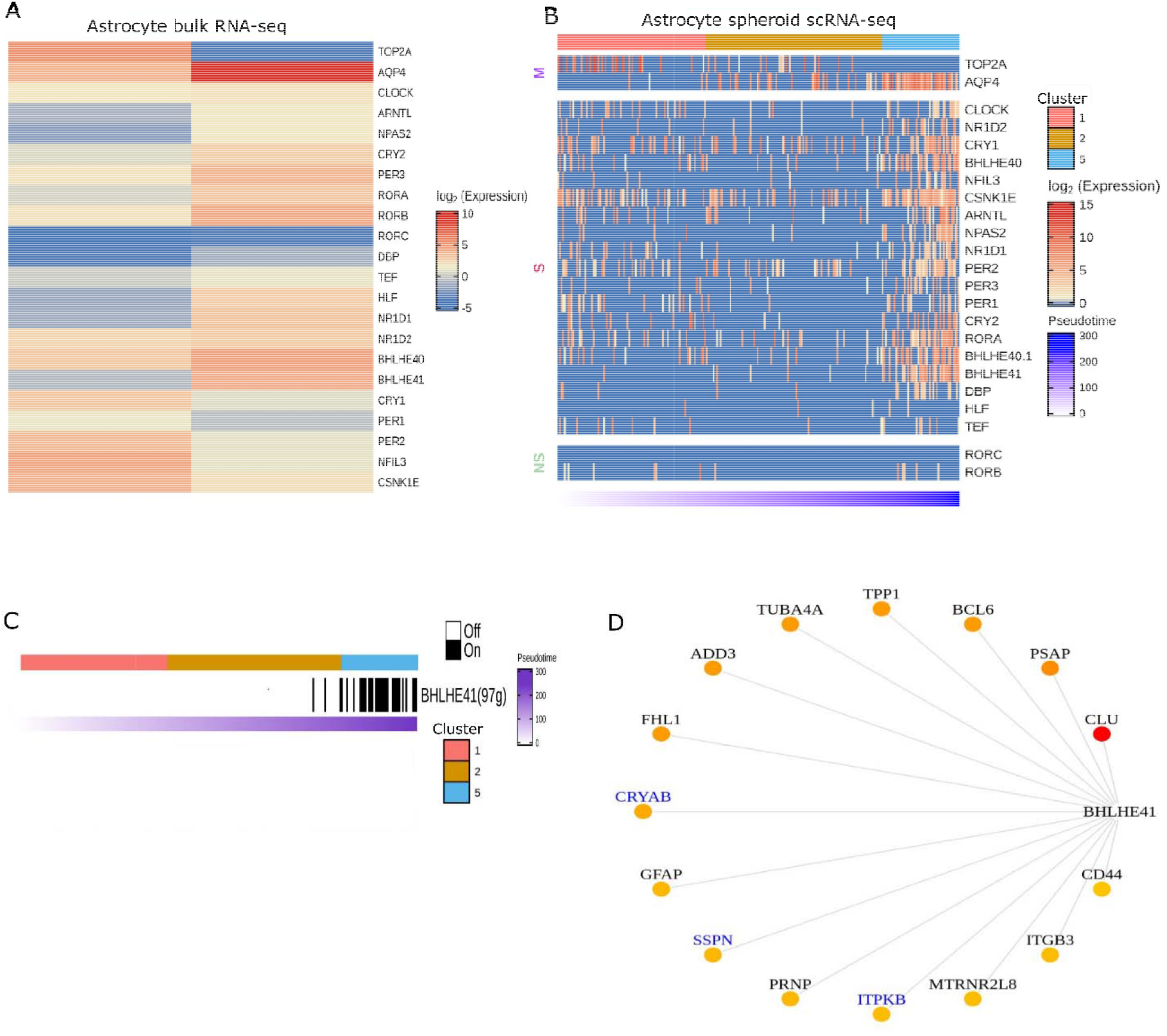
Expression dynamics of CGs-CNs during astrocyte genesis. A) Heatmap showing CGs expression in fetal and adult brain derived astrocytes profiled by total RNA-seq. TOP2A and AQP4 represent marker genes associated with early and late stages of astrocyte development respectively. B) Heatmap showing CGs (row) expression over pseudotime ordered cells (column) derived from astrocytic spheroid, M denoting marker genes, S and NS depicting genes showing statistically significant and non-significant pseudotime expression dynamics respectively. Top annotation denotes cell clusters assigned by TSCAN, bottom annotation denotes TSCAN calculated pseudotime in ascending order. C) Binary activity heatmap of CGs modules over pseudotime aligned trajectory of astrocyte genesis. Top annotation denotes cell clusters assigned by TSCAN, bottom annotation denotes TSCAN calculated pseudotime in ascending order. D) Network plot of BHLHE41 module in astrocytes, the edges are placed from highest (red) to lowest (yellow) importance score of target genes. GeneMANIA reported co-expression gene pairs are highlighted in blue color, only top15 target genes are shown.

To strengthen this observation, we analyzed single-cell transcriptomic data of astrocytes derived from cortical spheroids (Sloan et al. 2017). Similar to neuronal differentiation analysis, we first constructed pseudotime trajectory of astrocyte differentiation and classified cell clusters using cellular markers of early and late stages of astrocyte development. Overall, we found an increased expression of all CGs except RORB and RORC in differentiated astrocytes (Figure 4B). This was in concordance with the trend seen in total RNA-seq of fetal and adult brain derived astrocytes.

The co-expression analysis of astrocyte scRNA-seq revealed the BHLHE41 associated module with 97genes to be the prominent module which showed enrichment in the later stages of astrocyte genesis (Figure 4C). BHLHE41 acts as a transcriptional repressor in the circadian regulation of gene expression. Previous studies have reported association of mutations in BHLHE41 with short sleep timing in both humans and experimental models (He et al. 2009; Pellegrino et al. 2014). Also, it shows high abundance in astrocytes as compared to neurons (Y. Zhang et al. 2014). Few of the topmost target genes in the BHLHE41 module included CLU, PSAP, BCL6, CRYAB, TPP1, TUBA4A and ADD3 (Figure 4D). The co-expression of a few genes like ITPKB, SSPN and CRYAB have been previously reported in the GeneMANIA. Additionally, we report various novel genes, specifically GFAP, a well known marker of astrocytes has been shown to be associated with the etiology of Alexander disease (Li et al. 2018). We also observed CD44 co-expression with BHLHE41, CD44 acts as a vital adhesion protein during astrocyte development (Y. Liu et al. 2004) and an increase in CD44 positive astrocytes have been shown in Alzheimer’s brain (Akiyama et al. 1993). Our approach highlights the potential regulation of these vital genes involved in astrocyte function by BHLHE41.

## Discussion

We combined pseudotime/co-expression analysis and explored potential regulatory contributions of CGs in the developing brain. Our workflow reveals CGs-CNs with cell type and cell state identity, a feat not possible from bulk RNA-seq based co-expression analysis. Also, high gene drop-out rates pertaining to scRNA-seq greatly impact the pseudotime analysis of low expressing genes like CGs. Therefore, we looked at co-expression modules which can accurately reflect variations in gene expression trends related to underlying biological processes (neurogenesis).

Our analysis of the murine and human embryonic brain scRNA-seq datasets reveals CGs-CNs of RORB, whose expression and role in cortical development is well studied in murine models. Further, many of the identified co-expression pairs are also reported in GeneMANIA with few of the target genes showing oscillatory expression in the adult brain (Seney et al. 2019), again highlighting the biological relevance of our analysis.

Many recent studies reveal the regulatory role of non-neuronal cell types in brain physiology, our analysis also identified four CGs-CNs in the non-neuronal population of the developing human brain. Among the four modules, BHLHE41 genes strongly co-express with genes having crucial function in microglia such as P2RY12 and CLEC7A. Further, BHLHE41 upstream coexpression analysis shows TFs with known microglia function like TAL1, IRF8, IKZF1 and ZBTB24. In addition, astrocytes derived from cortical spheroids show high expression of BHLHE41 module in mature astrocytes. Above results clearly favor BHLHE41 as a prioritized module for future biological validation in the context of glial biology.

Our meta analysis study uncovers CGs-CNs in diverse cell types of the developing brain. We believe that with exponential increase in new scRNA-seq datasets, with improved sample preparations such as pre-sorting of rare cell types and better sequencing depth will further reveal novel CGs-CNs. In future, similar analysis approach can be extended to generate CGs-CN map of all cell types in the human body.

## Methods

### Pseudotime analysis

TSCAN was used for pseudotime analysis (https://github.com/zji90/TSCAN). Single-cell gene expression matrix was subjected to TSCAN default workflow (Figure 1A), the workflow included preprocessing, dimensionality reduction followed by clustering. The genes with zero count were removed and scRNA-seq dropouts were handled by clustering genes having similar expression patterns using euclidean distance and complete linkage. The above preprocessing method was applied to all the datasets. Next, PCA was used for dimension reduction followed by clustering (mclust), yielding minimum spanning tree (MST). The identification of clusters was performed using marker genes. p-value and FDR was calculated on the TSCAN ordered cells with an FDR of <0.05. TSCAN assigned cell clusters, were inspected for a panel of marker genes from the publications of the respective datasets, while a single marker gene was used for visualization purpose. If required, TSCAN cluster ordering was further manually corrected to place neural stem cell clusters as the starting point of the pseudotime.

The accession number of the datasets and their TSCAN path is provided in supplementary table 2 and 3 respectively.

### Co-expression analysis

Two steps in the pySCENIC (V0.10.3) (Aibar et al. 2017) pipeline were implemented in our co-expression analysis. First the co-expressing gene modules were identified using GRNBoost (https://github.com/aertslab/GRNBoost). Input for GRNBoost included a list of annotated transcription factors (https://github.com/aertslab/pySCENIC/tree/master/resources) and scRNA-seq expression matrix. In return GRNBoost outputs adjacency matrix with TFs, target genes and importance score. All the CGs (TFs) and its associated targets genes with importance scores greater than 99th percentile were subsetted as CGs-CNs. pySCENIC-GRNBoost was implemented through the command line in linux. All the original adjacency matrices are provided in supplementary data.

Next AUCell (V1.11.0) was used for scoring the activity of above CGs-CNs. AUCell uses “Area under curve” to score the activity of a given geneset (CGs-CNs) within expressed genes of scRNA-seq dataset. AUCell was implemented using R (4.0.2) using default parameters. The CGs-CNs which crossed the default threshold of AUCell geneset activity parameters are shown as binary CGs-CNs activity heatmap.

### ScRNA-seq dataset description

#### Mouse Cortex

Yuzwa et al. performed drop-seq of CD1 mouse embryos at E11.5, 13.5, 15.5 and 17.5 stage. For E15.5 embryonic stage, 13 embryos were used and their brains dissected and dissociated for Drop-seq droplet collection. Following droplet collection single-cell libraries were prepared and sequenced on Illumina NextSeq500 platform.

#### Mouse Subventricular zone (SVZ)

Babodilla et al performed microdissection of SVZ from C57BL/6 adult mice. GLAST^+^ Prominin1^+^ and PSA-NCAM^+^ were used as marker to sort neural stem cells and neuroblasts from SVZ region. scRNA-seq libraries were prepared using Smart-seq2 and sequenced on HiSeq2000 platform. Trimmed reads were mapped to the mouse genome (ENSEMBL Release 78) using STAR_2.4.0g followed by read quantification as FPKM.

#### Mouse Subgranular zone (SGZ)

Shin et al. used transgenic mice expressing nuclear localized CFP under the control of nestin Nes-CFP^nuc^). The CFP containing cells marking qNSCs and their progenies were sorted under a fluorescent microscope using glass pipette. The single-cell libraries were prepared by SMART protocol and sequenced by HiSeq2500 followed by read mapping and quantification to give TPM matrix.

### Developing human fetal cortex

Nowakowski et al. performed stereotaxic based microdissection of brain regions like prefrontal cortex (PFC), primary visual cortex (V1) and MGE, the brain region aged from 5-37pcw. The single-cell suspension was prepared using FluidigmC1 followed by sequencing of single-cell libraries on the Illumina Hiseq2500 platform. The sequencing reads were aligned to GRCh38 and quantified to reveal transcript count as CPM.

### Astrocyte spheroid

Astrocyte lineage cells were purified from cortical spheroids derived from human iPSC line by Sloan et al. The immunopanning method was used to isolate astrocytes using HepaCAM as an astrocyte marker. The sequencing libraries were prepared from astrocytes derived using immunopan method at 100, 130, 175 and 450 days of *in vitro* differentiation. Single-cell libraries were prepared using SMART-seq protocol and sequenced on NextSeq500 sequencing platform. The raw reads were aligned to hg19 human genome followed by transcript quantification as FPKM values.

### Total RNA-seq

#### Human astrocytes

Zhang et al. performed RNA-sequencing of astrocytes derived from juvenile (8-18years old) and adult (21-63 years old) human brain. The astrocytes from the brain tissue were purified in culture using immunopanning. Briefly, single-cell suspension of donor tissues was subjected to dishes coated with antibodies against various cell types, Hepa-CAM was used as a marker for astrocytes. The cDNA libraries from the two samples were sequenced on illumina NextSeq sequencer as 150bp paired end reads. The RNA-seq reads were mapped using TopHat2 to hg19 as the reference genome and expression level was estimated as FPKM.

## Acknowledgements

The authors acknowledge Dr. Beena Pillai and Dr. Souvik Maiti for their help in writing the manuscript and discussions. We are grateful to them for allowing us to communicate the manuscript. We also acknowledge the high performance computing (HPC) facility of CSIR-IGIB.

## Author contributions

SR and SS contributed equally. SR and SS conceptualized the work, SR and AHA performed the analysis, SR and SS prepared the manuscript.

## Supplementary files

TableS1: List of core clock genes analyzed in the study.

TableS2: Accession number of datasets used for the analysis.

TableS3: Details of TSCAN pseudotime clusters.

TableS4. Identities of cell clusters in developing human brain.

Table S5, S6, S7 and S8: GRNBoost derived adjacency matrix of mouse E15.5 cortex, mouse SVZ, developing human brain and astrocytes respectively.

Figure S1. Rorb co-expression network in mouse embryonic cortex E15.5.

Figure S2. Heatmap of pseudotime ordered cells in mouse SGZ.

